## Supplementary material for "Causes and consequences of experimental variation in *Nicotiana benthamiana* transient expression": Nicotiana Growth Protocols

**Full-length protocol**

This protocol has been optimized for the JBEI plant growth room conditions (long day, ~120 ppfd, 23°C, 65% humidity). Plant needs may vary when humidity / temperature / light intensity / light cycle are altered. All plants cycle through dedicated 1 week-, 2 week-, 3 week-, and 4 week-old zones to minimize batch-to-batch variation.

Soil prep: Sungro Sunshine mix #4 (aggregate plus), supplement with Osmocote (14-14-14) pellets at 1.5 TBSP (~20 mL) / 4L. Mix in the osmocote well and manually break apart large chunks of soil. When preparing soil for seedlings, a layer of Pro-Mix PGX is added as topsoil. Plants are grown in Greenhouse Megastore 3.11" x 3.11" x 2.25" pots (Traditional Insert: CN-IKN-1801) which fit 18 pots per flat (1020 Trays Heavy Duty: CN-FLHD-X2).

If tasks are split between multiple individuals, we recommend that the same individual(s) do the same tasks each week to minimize batch-to-batch variation.

***Day 0: Germinate seeds***

- Fill an adequate number of pots with osmocote-supplemented soil (usually ~50-100 good seedlings per pot).
- Wet the soil by pouring excess tap water from above and allowing it to drain through
- Add a liberal layer of topsoil, and thoroughly wet it. Since topsoil is very fine and tends to be hydrophobic, this is easiest to do with a spray bottle.
- Using a spatula, sprinkle a pinch of seeds over the soil, ideally 50-100 seedlings per pot. If they germinate too densely, they will be smaller at transplant time. More sparse, healthy seedlings tend to grow into larger, healthy mature plants.
- Pour 0.5 L of tap water into the bottom of the tray to ensure high humidity.
- Put pots into a tray and cover with a hood (all vents closed) to keep humidity high
- Seedlings should be ready to transplant in one week

***Day 7: Transplant seedlings***

- Prep soil with osmocote as described above. For each flat to be transplanted, take a sheet of 18 pots, place it in the flat tray, and fill all to the brim. (Flats can be stacked to conserve space).
- Break out only the corner pot from each flat and add 3 L of tap water to the bottom of the tray.
- Allow the soil to soak for at least 60 minutes, until the top of the soil looks wet.
- After soaking, pour the excess water out from the bottom of each tray.
- Break apart the 18 pots in the flat. If the plants get big and the pots are still connected, it is very difficult to separate them without damaging the plants.
- Transplant seedlings.
  - Make a small hole in the soil of all the seedling pots.
  - Gently remove seedlings from the germination pot and place them inside their own individual pot, root inside the hole.
  - Once all the seedlings are placed in their own pot, push the soil compact around the roots to secure the seedlings in the soil.
  - *Be careful not to damage the roots*. Any seedling with a snapped root will be stunted and may not survive transplantation; throw it away. Grip seedlings by one of the cotyledons (forceps may help) so as not to damage the stem/roots.
- Place a hood over each flat, *hood vents closed*, and move the flat into a growth room/chamber into the 1 week-old plant zone. The transplanted seedlings should require no attention for the next week.

***Day 14: Open hood vents***

- Move the post-transplantation flats from the 1 week-old plant zone to the 2 week-old plant zone. Open the top and both side vents on all of their hoods.
- This is important to allow the humidity within the hood to slowly equilibrate to the outside conditions. Removing the hood all at once causes a rapid humidity change which can be detrimental.

***Day 19: Remove hoods***

- Remove the hood from each flat.

***Day 21: Water plants***

- Move flats from the 2 week-old zone to the 3 week-old zone.
- Give each flat 1 L of tap water. It is easiest to pour the water into the bottom of the tray using a funnel.

***Day 26: Separate plants into 9/flat and water***

- Move the flats from the 3 week-old zone to the 4 week-old zone.
- Separate the 18 plants in each flat into two flats with 9 plants each, checkerboard pattern to maximize space for each plant.
- Checkerboarding helps prevent overcrowding, shade avoidance, and loss of structural integrity.
- Give each flat 1 L of tap water. Pour directly into the bottom of the tray.

***Day 29: Agroinfiltration***

- Infiltrate plants as desired.
- Post-infiltration, give each tray of plants 1 L of tap water so they are not dried out 72 hours later.

**Quick guide**

***Tuesday***

- Remove the hoods from flats in the 2 week-old zone.
- Separate flats in the 3 week-old zone into 9 plants/flat (checkerboarded). Move into the 4 week-old zone and give each flat 1 L of tap water.

***Thursday***

- Prepare soil for as many flats as desired and saturate each in 3 L water/flat for at least 1h.
- Move the flats in the 2 week-old zone to the 3 week-old zone. Add 1 L of tap water/flat.
- Move the flats in the 1 week-old zone to the 2 week-old zone. Open all vents on the hoods of these flats.
- Drain the water from the flats for transplantation and break apart the pots.
- Transplant one seedling to each pot, cover with a hood (all vents closed), and place in the 1 week-old zone.
- Sow seeds for next week.

***Friday***

- Infiltrate.
- Recover each flat of infiltrated plants in the 4 week-old zone with 1 L of tap water/flat.

**Example calendar**

Letters represent unique plant batch IDs. IDs are in alphabetical order from oldest to youngest.

| **Wk** | **Mon** | **Tue** | **Wed** | **Thu** | **Fri** |
| --- | --- | --- | --- | --- | --- |
| 1 |  |  |  | Sow seeds (A) |  |
| 2 |  |  |  | Sow seeds (B)  Transplant seedlings (A) |  |
| 3 |  |  |  | Sow seeds (C)  Transplant seedlings (B)  Open vents (A) |  |
| 4 |  | Remove hoods (A) |  | Sow seeds (D)  Transplant seedlings (C)  Open vents (B)  Water (A) |  |
| 5 |  | Remove hoods (B)  Separate pots, water (A) |  | Sow seeds (E)  Transplant seedlings (D)  Open vents (C)  Water (B) | Infiltrate, water (A) |
| et cetera | | | | | |
