## Supplementary material for "Causes and consequences of experimental variation in *Nicotiana benthamiana* transient expression": Modelling

**Monte Carlo Simulation**

A dataset of 1,813 *N. benthamiana* plants from 32 independent GFP transient expression experiments was compiled, spanning multiple years, researchers, and binary vector designs using either *A. tumefaciens* GV3101 (1,087 plants) or EHA105 (726 plants). This dataset captures the real-world variability of transient expression performance outside of a single controlled experimental environment.

For each plant, a coefficient of variation (CV) of GFP expression was calculated from eight leaf disks. Using these data, we parameterized a hierarchical Monte Carlo model to reproduce observed variability at three levels: (i) week-to-week shifts in average plant CV, representing overall plant quality between experiments; (ii) plant-to-plant heterogeneity within a given week; and (iii) within-plant disk-level noise.

Weekly median plant CVs followed a log-normal distribution with parameters meanlog = -1.447, sdlog = 0.300 for GV3101 and meanlog = -0.903, sdlog = 0.460 for EHA105. The within-week spread of plant CVs (weekly SD-of-CV) also followed a log-normal distribution (meanlog = -2.3248, sdlog = 0.3897). Fit adequacy was confirmed using Kolmogorov–Smirnov and density RMSE metrics.

These data were used to simulate variance across random weeks. For each simulated experiment, a week quality value was selected using a random number from the log-normal distribution of the strain of choice (GV3101 vs EHA105), representing the average plant CV for that week. Additionally, a week quality spread value—modeled from the empirical standard deviation of plant CVs across different experiments—was randomly selected from the underlying log-normal distribution and used for the standard deviation of CV values for that week. These two values were then used to generate individual plant CVs for the simulated experiment, incorporating both the mean and standard deviation values for plant CV that were randomly generated for that week. Each plant drawn from this distribution thus has its own unique CV value, bounded by biologically relevant values between 0.03-2.0, which were inferred from the 1813 measured plant spread.

For each simulated plant, 8 leaf discs were generated using an arbitrary mean value of 100,000 GFP units and standard deviation derived from the plant CV. A given simulated experiment would then have all leaf disc data bulked (e.g. 10 plants with 80 total discs) and used for population-level comparisons. For the modeling in this study to determine features such as minimal detectable effect size or power analyses, 1000 weeks of experiments were simulated using the above pipeline. Simulation outputs were summarized as statistical power (fraction of iterations with significant construct differences) across plant counts and effect sizes. Comparison of features such as effect size was conducted through two-sided t-tests with Benjamini–Hochberg FDR control at α=0.05. For power analysis modeling, a ≥95% accurate detection of differences was used as a threshold for classifying sufficient detection power.

To validate the outputs of this simulation, goodness-of-fit between simulated and empirical CV distributions was assessed using KS and 1D Wasserstein distances (Fig. 6b).

**Mixed effect modeling**

Modeling of variance was conducted using the pCM2::GFP dataset which consisted of GFP expression data from 114 plants collected over 15 weeks. Fluorescence data (n = 8 discs per plant) were transformed to a log10 scale to linearize multiplicative effects and to account for heteroscedasticity.

On this scale, a linear mixed-effects model was fit to account for 4 major sources of variation: (i) experiment-to-experiment shifts in plant mean, (ii) experiment-to-experiment shifts in intraplant variance, (iii) plant-level shifts in mean GFP expression, and (iv) leaf-to-leaf expression variability, along with a residual component. The equation for modeling such variance for the GFP expression of a given leaf disc is as follows:

**Log_10_GFP(_disc, plant, week_) =**

**β0** (dataset mean) +

**β1 * Leaf position** (dataset difference between top and bottom leaf) +

**u_0_** (week’s deviation in mean GFP) +

**u_1_ * Leaf position** (weeks deviation in top vs bottom leaf difference) +

**v_0_** (Individual plant’s mean deviation) +

**v_1_** (Individual plant’s leaf-to-leaf variation) +

**e** (residual error)

This was calculated using the R code:

m_final <- lmer(

log_GFP ~ Leaf_c +

(1 + Leaf_c | Date) +

(1 + Leaf_c | Unique_plant_ID),

data = PC4_data, REML = TRUE)

In the above, Leaf_c encoded the contrast between the top and bottom leaves.

**Disc-to-disc, positional, and technical variance:**

To determine the impact of disc-to-disc variability, leaf position, and technical replication noise on total observed variability, the same pCM2::GFP construct was used to infiltrate 24 plants on the same two leaves (T4 and T5) in two positions (proximal to the petiole and distal) with two *Agrobacterium* strains (GV3101 and EHA105, 12 plants per strain). For each leaf, eight discs were taken (four per proximal or distal position). GFP fluorescence was quantified as before, except plates were scanned three times each in their standard position and then were flipped 180 degrees and scanned three more times to enable quantification of technical variance. The following R code when then used to fit a mixed effects model to better account for residual variation:

m_leafc <- lmer(

log_GFP ~

Plate_Position + (location of disc on the plate) +

Leaf_c + (Leaf T4 or T5 variance)

(1 + Leaf_c | Plant) + (Plant mean and leaf-to-leaf variance)

(1 | Plant:Leaf:Site) + (Proximal vs distal position impact)

(1 | DiskID) + (disc-to-disc variation),

data = punchvar, REML = TRUE)

**Ruby and PDC modeling**

Modeling of metabolite production variance was conducted in a similar way but with reduced variables given tissue bulking that was done prior to extraction. For Ruby, a single infiltrated spot per leaf was excised and extracted, resulting in data for only week-to-week and plant-to-plant variability. As only a single measurement was taken per leaf, the residual of this model includes leaf-to-leaf variability, as calculated by the following R code:

m_ruby <- lmer(

log10_ruby ~ 1+

(1 | Experimental_replicate) + (experiment mean variation)

(1 | Experimental_replicate:Plant), (plant level variation within experiment)

REML = TRUE)

For PDC production, tissue from both leaves was bulked prior to extraction. For this reason, a linear model was used to examine the relationship between week and PDC production, with residual variance representing plant-to-plant and leaf-to-leaf variability not explained by week-to-week differences.
