## Supplementary Figures for "Causes and consequences of experimental variation in *Nicotiana benthamiana* transient expression"

### **Supplementary information**

| **Strain** | **Description** | **Reference/Source** |
| --- | --- | --- |
| *E. coli* XL1-Blue | Cloning strain of *E. coli* | Agilent |
| *Agrobacterium tumefaciens* GV3101:pMP90 | Laboratory strain of *A. tumefaciens* C58 that has been disarmed of T-DNA | Intact Genomics |
| *Agrobacterium tumefaciens* EHA105 | Laboratory strain of *A. tumefaciens* C58 with a hypervirulent pTiBo542 Ti plasmid that has been disarmed of T-DNA | GoldBio |

| **#** | **Origin** | **T-DNA description** | **Resistance marker** | **Part ID** |
| --- | --- | --- | --- | --- |
| 1 | pVS1 | PCM2:GFP_CaMV35S2:nptII | kanamycin | JPUB_021048 |
| 2 | pVS1 | PCH5:CYP76AD1 | kanamycin | JPUB_026773 |
| 3 | pVS1 | PCH5:DODA | kanamycin | JPUB_026771 |
| 4 | pVS1 | PCH5:glycosyltransferase | kanamycin | JPUB_026775 |
| 5 | pVS1 | PCH5:RUBY | kanamycin | JPUB_026785 |
| 6 | pVS1 | 35S:LigA | kanamycin | JPUB_018689 |
| 7 | pVS1 | 35S:LigB | kanamycin | JPUB_018691 |
| 8 | pVS1 | 35S:LigC | kanamycin | JPUB_018693 |
| 9 | pVS1 | 35S:QsuB | kanamycin | JPUB_018707 |
| 10 | pVS1 | 35S:AroG | kanamycin | JPUB_018685 |
| 11 | pVS1 | PCM2:GFP | spectinomycin | JPUB_026733 |
| 12 | pVS1 | PCM2:mCherry | spectinomycin | JPUB_026735 |
| 13 | pVS1 | PCM2:GFP_PCM2:mCherry | spectinomycin | JPUB_026737 |
| 14 | pVS1 | PCM2:mCherry_PCM2:GFP | spectinomycin | JPUB_026739 |
| 15 | pVS1 | PCM2:GFP_mCherry:PCM2 | spectinomycin | JPUB_026741 |
| 16 | pVS1 | PCM2:mCherry_GFP:PCM2 | spectinomycin | JPUB_026731 |
| 17 | pVS1 | GFP:PCM2_mCherry:PCM2 | spectinomycin | JPUB_026743 |
| 18 | pVS1 | mCherry:PCM2_GFP:PCM2 | spectinomycin | JPUB_026745 |
| 19 | pVS1 | GFP:PCM2_PCM2:mCherry | spectinomycin | JPUB_026747 |
| 20 | pVS1 | mCherry:PCM2_PCM2:GFP | spectinomycin | JPUB_026749 |
| 21 | BBR1 | PCM2:mCherry | kanamycin | JPUB_026779 |
| 22 | BBR1 | PCM2:GFP | kanamycin | JPUB_026777 |
| 23 | pSa | PCM2:mCherry | kanamycin | JPUB_026783 |
| 24 | pSa | PCM2:GFP | kanamycin | JPUB_026781 |
| 25 | pVS1 | PCM2:GFP_REVTOCS_PCM2:mCherry | spectinomycin | JPUB_026765 |
| 26 | pVS1 | PCM2:mCherry_REVTOCS_PCM2:GFP | spectinomycin | JPUB_026767 |
| 27 | pVS1 | PCM2:GFP_REVTOCS_mCherry:PCM2 | spectinomycin | JPUB_026769 |
| 28 | pVS1 | PCL2:GFP | spectinomycin | JPUB_026751 |
| 29 | pVS1 | PCL2:mCherry | spectinomycin | JPUB_026753 |
| 30 | pVS1 | PCH5:GFP | spectinomycin | JPUB_026759 |
| 31 | pVS1 | PCH5:mCherry | spectinomycin | JPUB_026761 |
| 32 | pVS1 | PCL1:mCherry | spectinomycin | JPUB_026763 |
| 33 | pVS1 | PCM1:mCherry | spectinomycin | JPUB_026755 |
| 34 | pVS1 | PCH4:mCherry | spectinomycin | JPUB_026757 |

**Supplementary Table 1.** All binary vectors used in this study. T-DNA descriptions are written (left border)-(T-DNA)-(right border). Underscores indicate contiguous expression cassettes that are contained within the same T-DNA. Gene:promoter is used to indicate that the cassette is read in the opposite direction of promoter:gene. All expression cassettes are terminated by T_AtUbq3 with the exception of the nptII cassette in binary vector 1, which is followed by the CaMV 3’ UTR, and the RUBY (vectors 2-5) and PDC (vectors 6-10) binary vectors, which are followed by the CpMV 3’ UTR. *A. tumefaciens* strain GV3101 is resistant to rifampicin and gentamicin, and *A. tumefaciens* strain EHA105 is resistant to rifampicin.

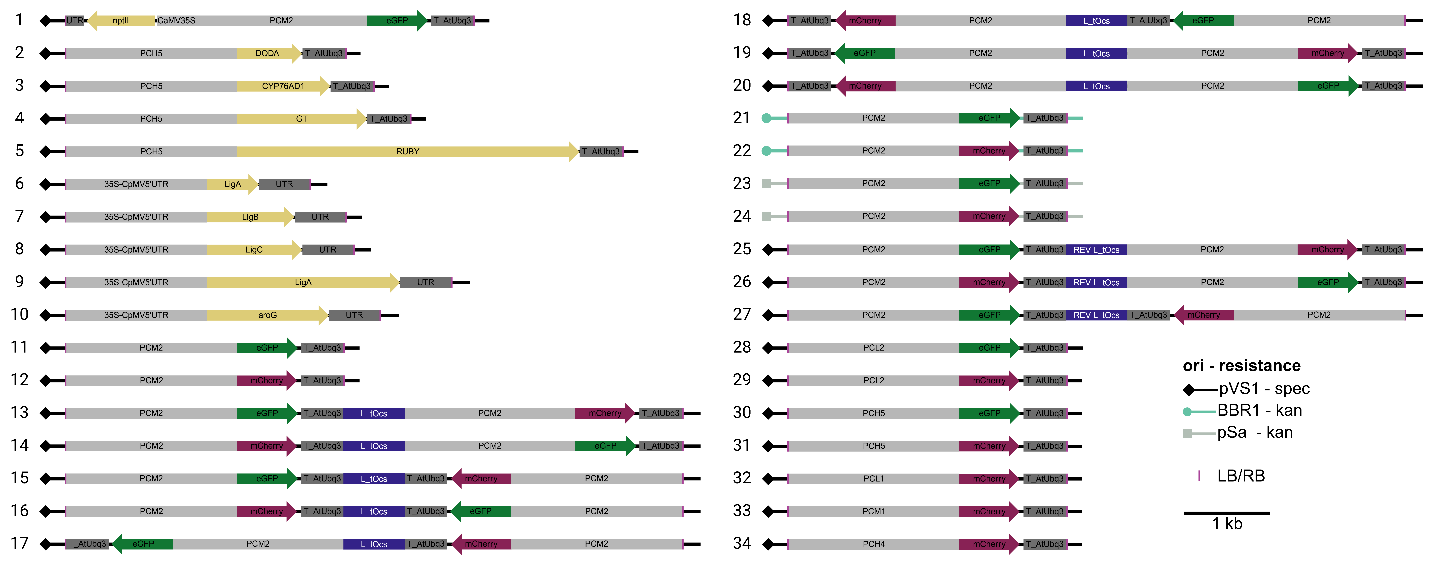
**Supplementary Figure S1.** Diagram of all binary vectors used in this study. Numbering of plasmids follows the numbering in Supplementary Table 1, *i.e.*, in order of appearance. T-DNAs are written from left border to right border. UTR in vector 1 is the CaMV 3’ UTR, and the UTR in vectors 2-10 is the CpMV 3’ UTR

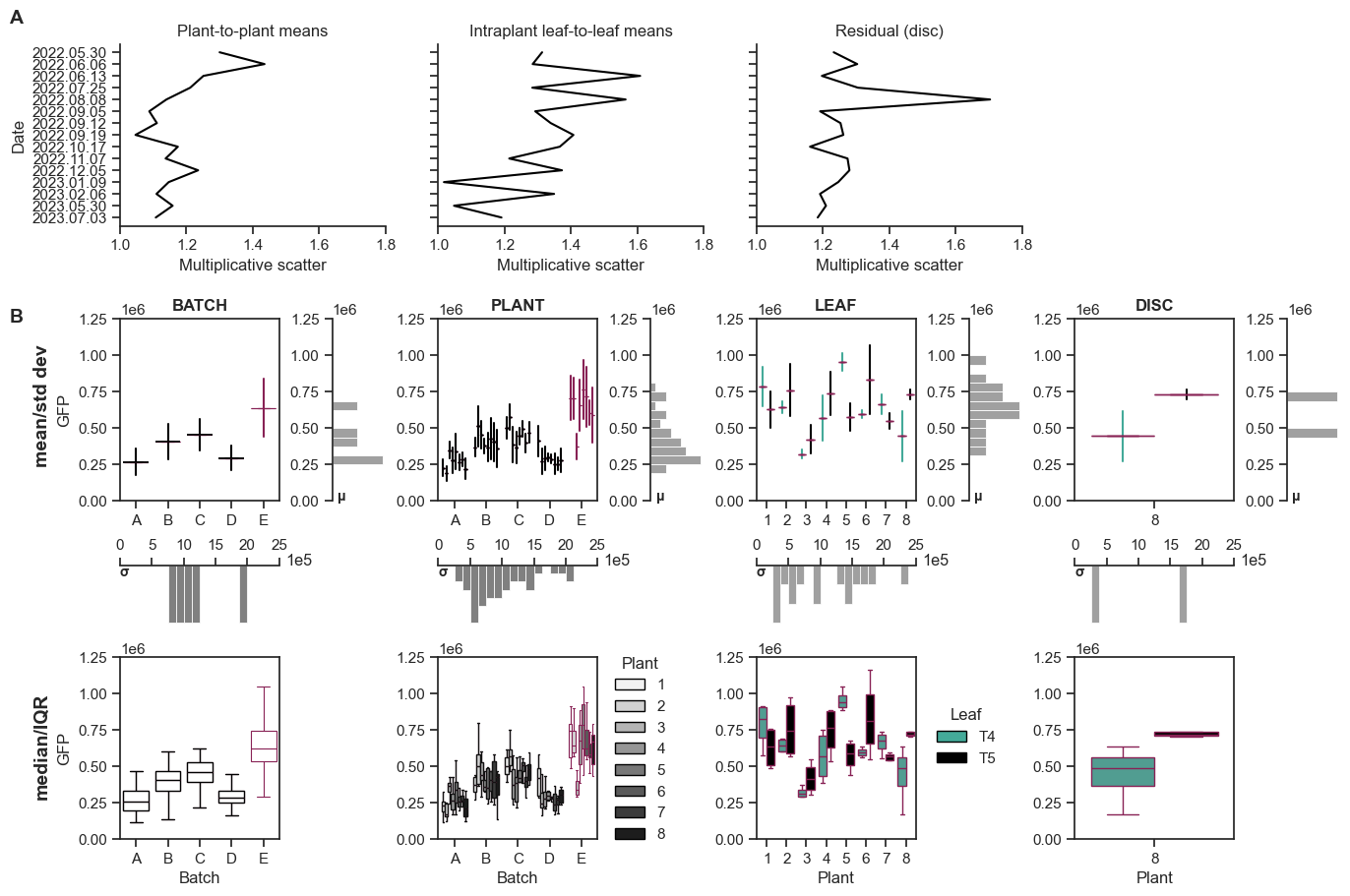
**Supplementary Figure S2.** Components of the mixed effect model. **A**, multiplicative scatters for different components of variance for 15 PCM2:GFP experimental replicates from Fig. 1. Multiplicative scatter indicates the factor by which the spread of a distribution around the mean is increased by a particular component of the variance, if all other factors are held constant. Multiplicative scatter of 1 indicates that the component contributes no additional variance. **B**, illustration of the components of the mixed effects model for five selected experimental replicates from Fig. 1. In the “mean/std dev” row, horizontal lines indicate the mean GFP and vertical lines are one standard deviation above and below the mean for a given batch, plant, or leaf. To the right of each subplot is a histogram of GFP means. *The mixed effect model treats the batch-, plant-, and leaf-level GFP means and standard deviations as having random intercepts and random slopes.* In the “median/IQR” row, the center and spread of the data are presented as boxplots (box is the IQR, middle line is the median, whiskers are the furthest point within 1.5*IQR).

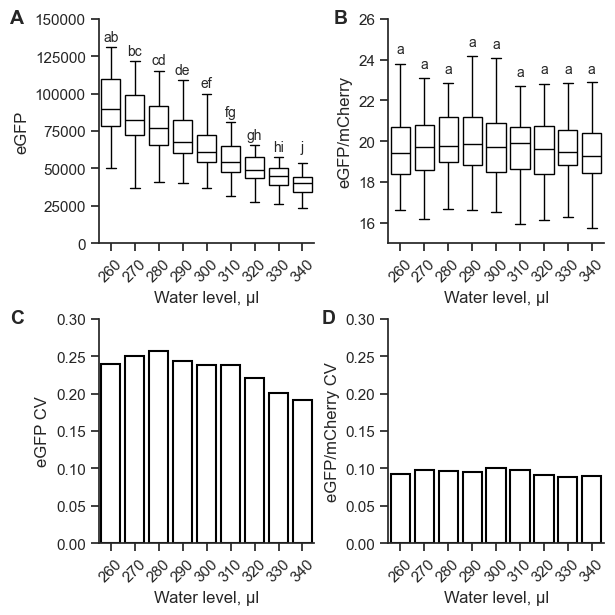

**Supplementary Figure S3.** Water level (in μL) effects on fluorescence signal strength and variability. 4-week old *N. benthamiana* plants were co-infiltrated using the same two *A. tumefaciens* GV3101 strains, each at an OD of 0.1 and carrying binary vectors with PCM2:eGFP:T_AtUbq3 or PCM2:mCherry:T_AtUbq3 inside the T-DNA.The volume of water was varied inside the 96-well plate used to measure the same leaf discs’ fluorescences. 8 plants, 2 leaves per plant, 4 discs per leaf. **A**, raw GFP fluorescence. **B**, ratio of GFP fluorescence to mCherry fluorescence. **C**, coefficient of variation of GFP fluorescence. **D**, coefficient of variation of GFP/mCherry fluorescence ratio. An independent 2-sample Student’s t-test and a Bonferroni correction were performed between every condition for both raw GFP fluorescence and for GFP/mCherry fluorescence ratio. p<0.05 was the cutoff for statistical significance. Shared letters indicate two conditions which are not significantly different from one another.

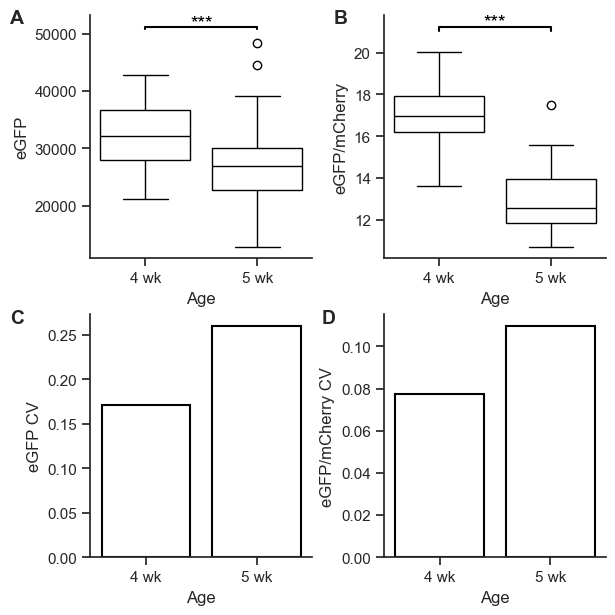

**Supplementary Figure S4.** Plant age effects on fluorescence signal strength and variability. *N. benthamiana* plants either 4- or 5-weeks old were co-infiltrated using the same two *A. tumefaciens* GV3101 strains, each at an OD of 0.1 and carrying binary vectors with PCM2:eGFP:T_AtUbq3 or PCM2:mCherry:T_AtUbq3 inside the T-DNA. 8 plants per condition, 2 leaves per plant, 4 discs per leaf. **A**, raw GFP fluorescence. **B**, ratio of GFP fluorescence to mCherry fluorescence. **C**, coefficient of variation of GFP fluorescence. **D**, coefficient of variation of GFP/mCherry ratio. An independent 2-sample Student’s t-test was performed between the 4- and 5-week plants. *** indicates p<0.001. Circles indicate outlier values beyond 1.5 times the interquartile range from the first and third quartiles.

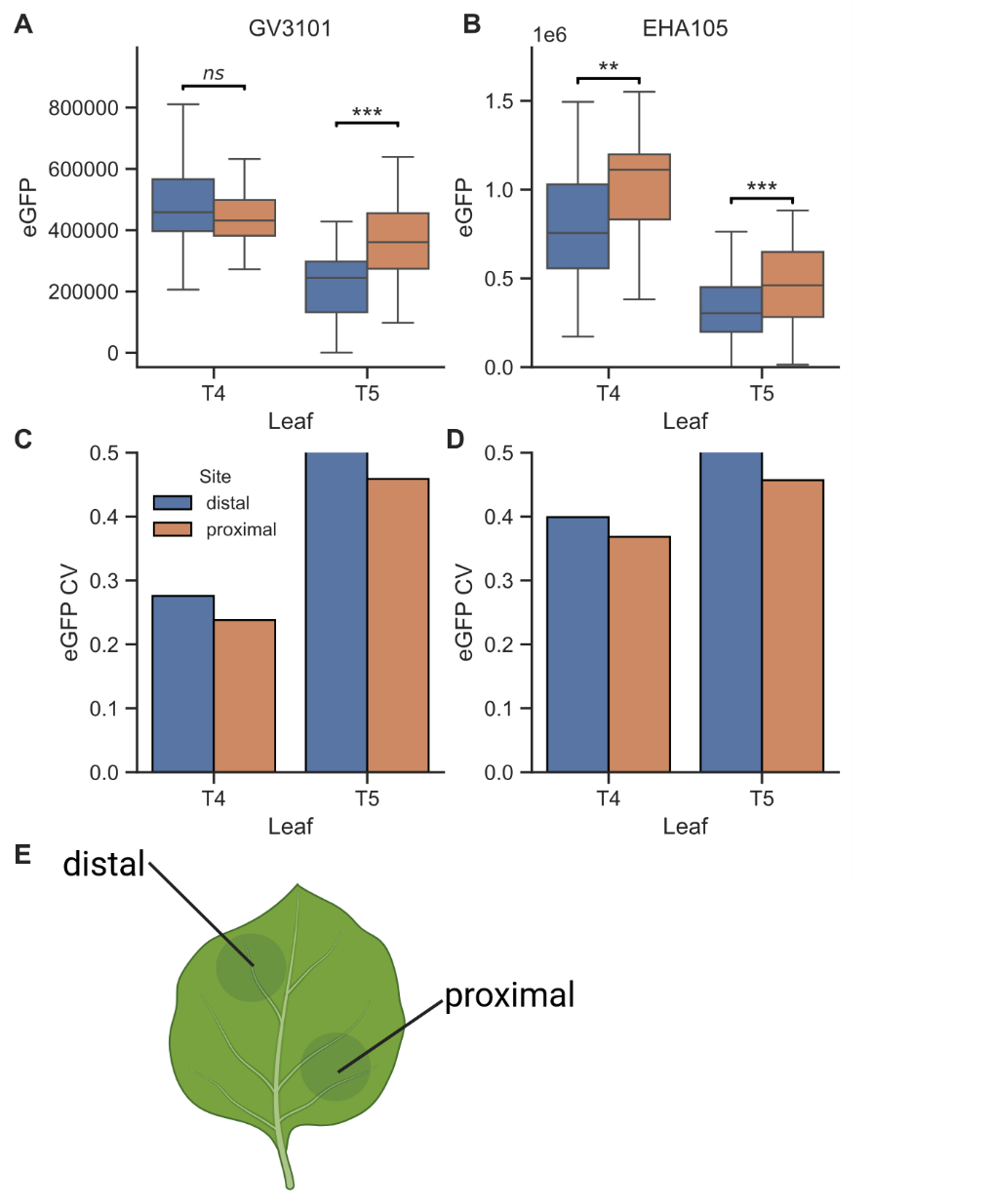

**Supplementary Figure S5.** Effect of leaf infiltration site on fluorescence signal strength and variability. eGFP measure in leaves T4 and T5 of 4-week old *N. benthamiana* plants infiltrated with either **A**, GV3101, or **B**, EHA105 carrying the same binary vector. Leaves were infiltrated at two sites, distal and proximal to the petiole. eGFP CV of **C**, GV3101 and **D**, EHA105. **E**, illustration of the proximal and distal infiltration sites. 12 plants per condition, 2 leaves per plant, 2 sites per leaf, 4 discs per site. A Student’s t-test was conducted to compare the distal and proximal sites for each leaf for each strain. No points are shown beyond 1.5 times the interquartile range.

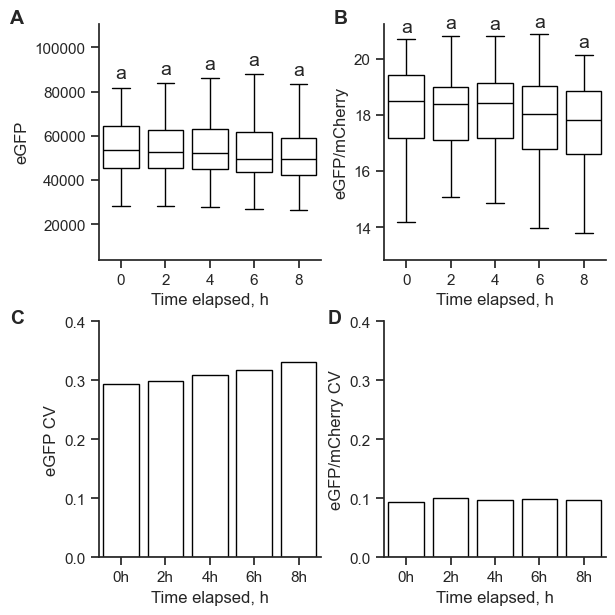

**Supplementary Figure S6.** Time elapsed between disc collection and measurement effects on fluorescence signal strength and variability. 4-week old *N. benthamiana* plants were co-infiltrated using the same two *A. tumefaciens* GV3101 strains, each at an OD of 0.1 and carrying binary vectors with PCM2:eGFP:T_AtUbq3 or PCM2:mCherry:T_AtUbq3 inside the T-DNA. 6 plants, 2 leaves per plant, 4 discs per leaf. Discs were collected and then measured on a plate reader every 2 hours thereafter until the end of the experiment. **A**, raw GFP fluorescence. **B**, ratio of GFP fluorescence to mCherry fluorescence. **C**, coefficient of variation of GFP fluorescence. **D**, coefficient of variation of GFP/mCherry ratio. An independent 2-sample Student’s t-test and a Bonferroni correction were performed between every condition for both raw GFP fluorescence and for GFP/mCherry fluorescence ratio. No statistically significant differences were observed with a p-value cutoff of 0.05. No points are shown beyond 1.5 times the interquartile range.

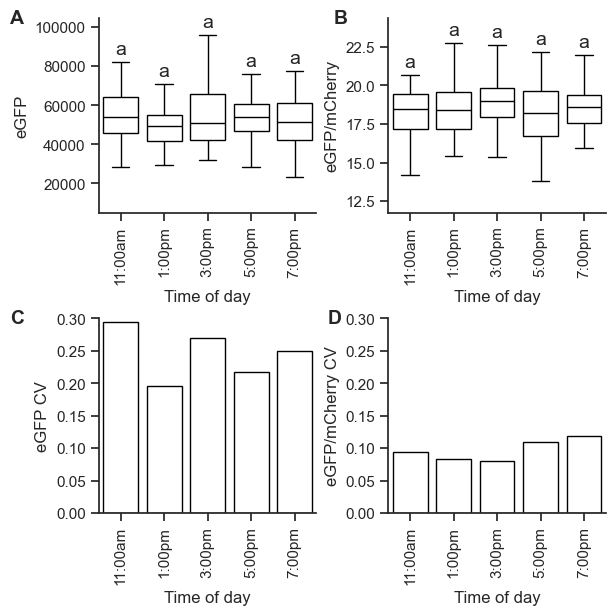

**Supplementary Figure S7.** Time of day during disc collection and measurement effects on fluorescence signal strength and variability. 4-week old *N. benthamiana* plants were co-infiltrated using the same two *A. tumefaciens* GV3101 strains, each at an OD of 0.1 and carrying binary vectors with PCM2:eGFP:T_AtUbq3 or PCM2:mCherry:T_AtUbq3 inside the T-DNA. 6 plants per condition, 2 leaves per plant, 4 discs per leaf. Each condition represents a group of plants from which discs were collected at the same time (every two hours from 11:00am to 7:00pm). Leaf discs were collected and immediately measured. **A**, raw GFP fluorescence. **B**, ratio of GFP fluorescence to mCherry fluorescence. **C**, coefficient of variation of GFP fluorescence. **D**, coefficient of variation of GFP/mCherry ratio. An independent 2-sample Student’s t-test and a Bonferroni correction were performed between every condition for both raw GFP fluorescence and for GFP/mCherry fluorescence ratio. No statistically significant differences were observed with a p-value cutoff of 0.05. No points are shown beyond 1.5 times the interquartile range.

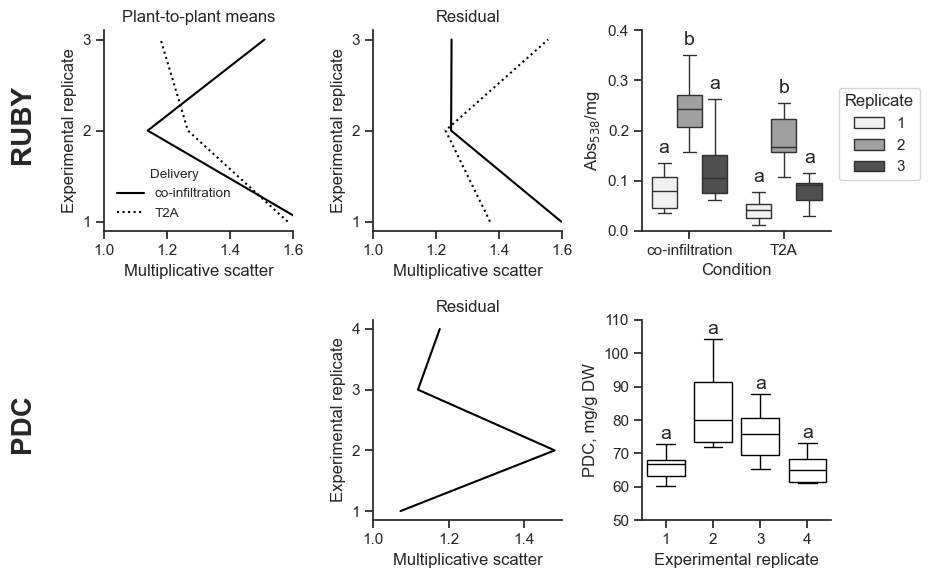

**Supplementary Figure S8.** Top row, left and center, multiplicative scatters for components of variation in betalain absorbance. Top row, right, absorbance at λ = 538 nm per mg dried tissuefor RUBY reporter agroinfiltrated into *N. benthamiana*. The three enzymes were introduced either on three separate T-DNAs (co-infiltration) or in one T-DNA connected by self-cleaving T2A peptides (T2A). Each independent experimental replicate was conducted with a unique plant batch on separate dates but extracted and measured together. 3 experimental replicates of n=6 plants, 2 leaves per plant, 2 co-delivery methods per leaf. Bottom row, center, multiplicative scatter for residual variance in 2-pyrone-4,6-dicarboxylic acid (PDC) yields. Bottom row, right, yields of the PDC biosynthetic pathway agroinfiltrated into *N. benthamiana*. Each independent experimental replicate was conducted with a unique plant batch on separate dates but extracted and quantified with HPLC together. 4 experimental replicates of n=6 plants, 2 leaves pooled per plant.

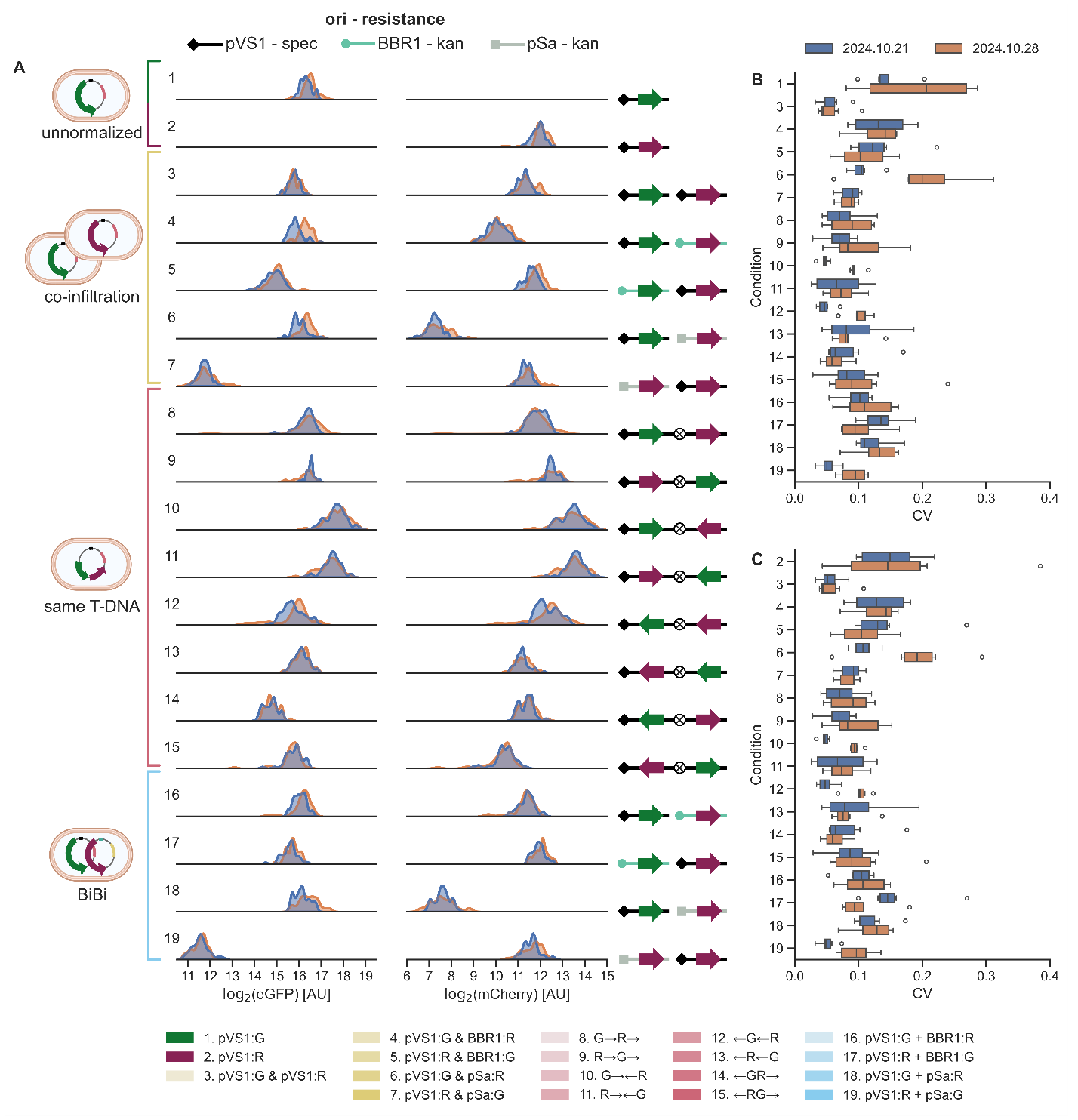

**Supplementary Figure S9.** Fig 2. data split by experimental replicate. Blue, 2024.10.21. Orange, 2024.10.28. **A)** Left, categories of delivery methods: unnormalized (green or magenta), co-infiltration (yellow), same T-DNA (pink), and BiBi (blue). Center, distributions of eGFP and mCherry fluorescence, n=48 leaf discs per experimental replicate per scheme. Right, cartoons showing the binary vector origin of replication, resistance marker, and orientations of FP expression cassettes in the T-DNA. All binary vector cartoons are read from left to right: ori, left border, T-DNA, right border. Origins of replication are pVS1 (diamond, black), BBR1 (circle, teal), and pSa (square, gray). Circles enclosing an X represent tOcs, a 722bp spacer in between the two expression cassettes. **B)** Plant coefficients of variation (CV), as calculated from the 8 discs per plant, when eGFP is treated as the reporter. All values are eGFP/mCherry CV except for scheme 1, which is GFP CV. **C)** Plant CVs when mCherry is treated as the reporter. All values are mCherry/eGFP CV except for scheme 2, which is mCherry CV. Normalization scheme IDs match across all subpanels. Total OD infiltrated in all schemes is 0.5. ODs of co-infiltrated strains are 0.25 each. In the legend, “&” indicates co-infiltration, arrows indicate the direction of an expression cassette, “+” indicates BiBi, and GFP is abbreviated to “G” and mCherry to “R”. Circles indicate outlier values beyond 1.5 times the interquartile range from the first and third quartiles.

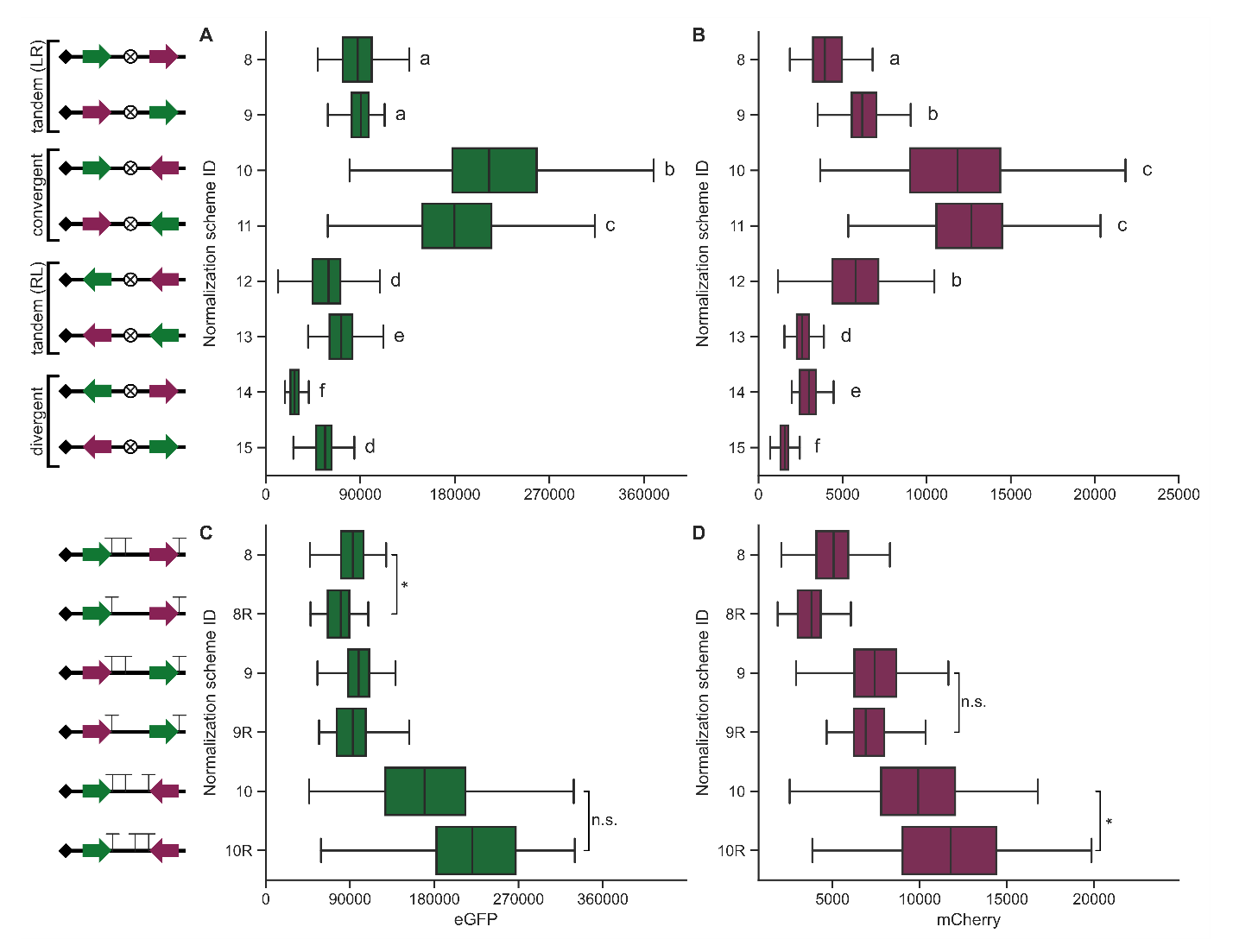

**Supplementary Figure S10.** **Impact of transgene orientation on gene expression.** Raw fluorescence signal from **A)** eGFP and **B)** mCherry for schemes 8-15 (two-cassette T-DNA schemes) from Fig. 2A. A two-tailed, independent Student’s t-test and Bonferroni correction were conducted between all pairs of schemes, and schemes with significantly different fluorescences are marked with different letters. The label “tandem (LR)” indicates left to right, and “tandem (RL)” indicates right to left. Two experimental replicates were performed for a total of n=96 leaf discs. **C)** eGFP and **D)** mCherry signals for schemes 8, 9, and 10 were also compared to another scheme where the spacer sequence tOcs was reversed (8R, 9R, and 10R). Reversal of tOcs converts a double terminator into a single terminator for schemes 8 and 9 and switches the doubly terminated cassette in scheme 10. Single terminators are represented as “T”, and double terminators are represented as “TT”. A one-tailed Student’s t-test was conducted to test whether the doubly terminated scheme has higher mean fluorescence than the corresponding singly terminated scheme. * indicates p<0.05. Six plants per condition, 2 leaves per plant, 4 discs per leaf. Binary vector cartoons are from left to right: ori, left border, cassette 1, 722 bp spacer, cassette 2, right border.

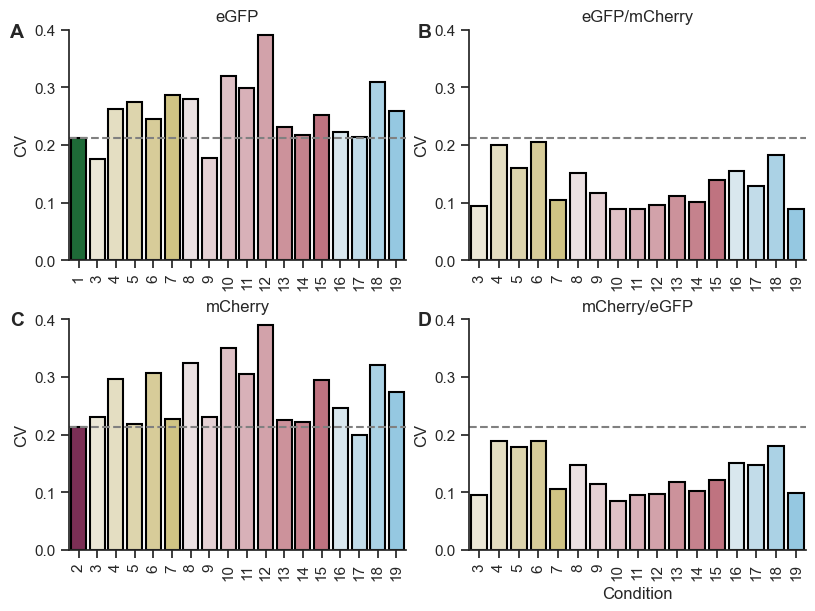

**Supplementary Figure S11.** Global CVs calculated from all discs from both experimental replicates, pooled, for **A)** eGFP, **B)** eGFP/mCherry, **C)** mCherry, and **D)** mCherry/eGFP of the normalization schemes from Fig. 2 (n=96 leaf discs). The dotted line in **A)** and **B)** is the eGFP CV of scheme 1, and the dotted line in **C)** and **D)** is the mCherry CV of scheme 2.

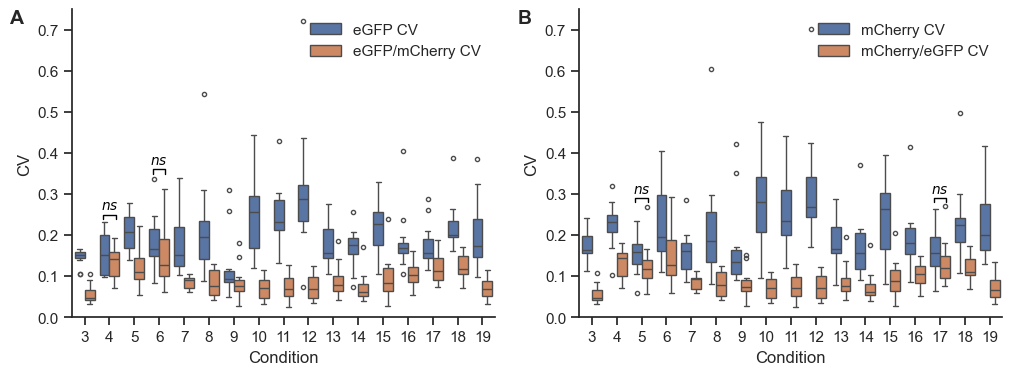

**Supplementary Figure S12.** Comparison of unnormalized CVs to normalized CVs from Fig. 2 when **A)** eGFP is the reporter and mCherry the normalizer or when **B)** mCherry is the reporter and eGFP the normalizer. A paired, one-tailed Student’s *t*-test was conducted, where each pair was the reporter CV and the reporter/normalizer CV from the same plant. The CV for each individual plant is calculated from 8 leaf discs. For visual clarity, only conditions for which the reporter/normalizer CVs were not significantly lower than the reporter CVs (p>0.05) are marked. Otherwise the reporter/normalizer CVs were significantly lower (p<0.05). Circles indicate outlier values beyond 1.5 times the interquartile range from the first and third quartiles.

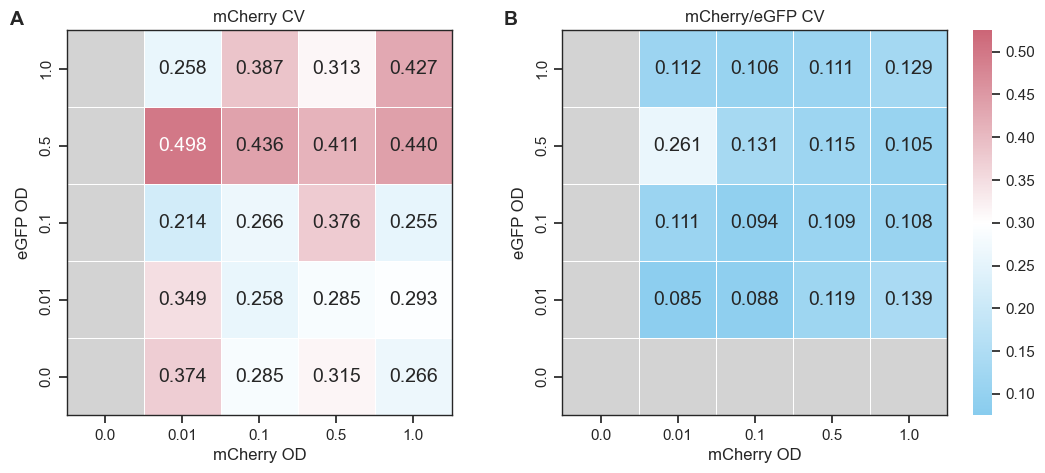

**Supplementary Figure S13.** Data from Fig. 3 showing mCherry as the reporter and eGFP as the normalizer instead. Matrices of all OD combinations’ CV of **A)** the mCherry fluorescence and **B)** ratio of mCherry/eGFP.

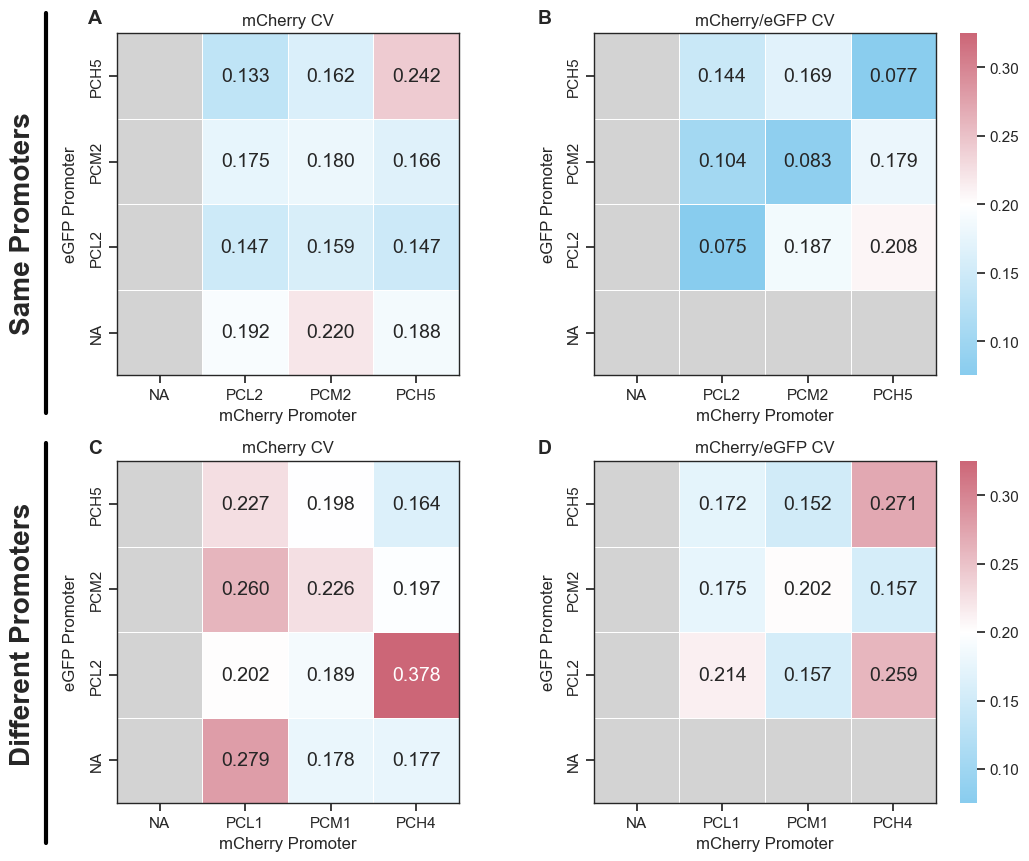

**Supplementary Figure S14.** Data from Fig. 4 showing mCherry as the reporter and eGFP as the normalizer instead. Matrices of all promoter combinations’ CV of **A)** the mCherry fluorescence and **B)** ratio of mCherry/eGFP when the set of 3 promoters are the same for eGFP and mCherry binary vectors (PCL2, PCM2, PCH5). Matrices of all promoter combinations’ CV of **C)** the mCherry fluorescence and **D)** ratio of mCherry/eGFP when the set of 3 promoters for mCherry binary vectors (PCL1, PCM1, PCH4) are different than the set for eGFP.

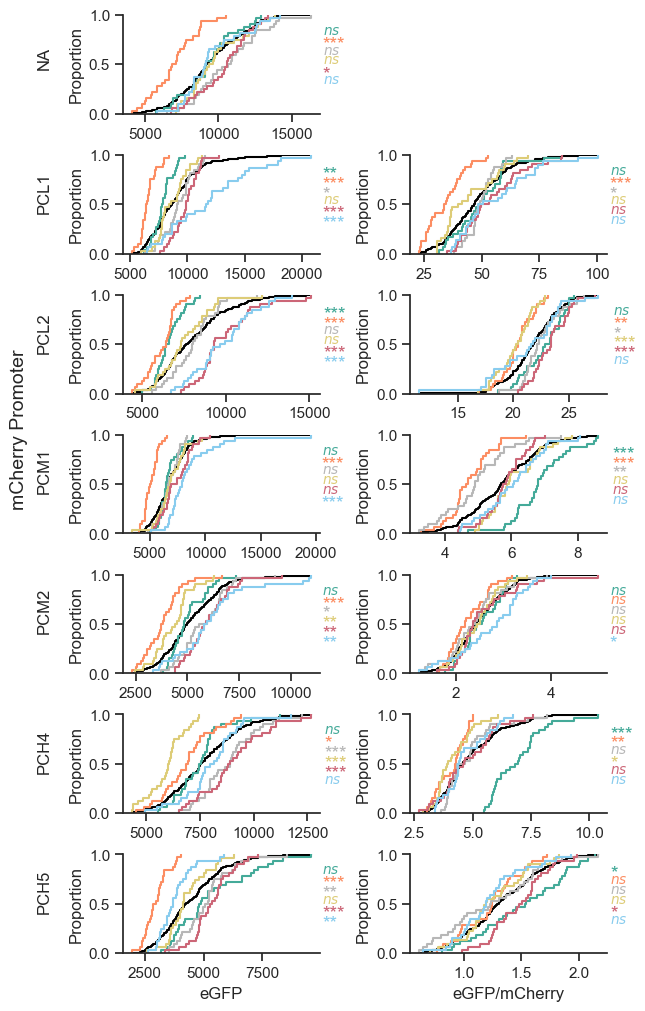

**Supplementary Figure S15.** Cumulative density functions of all conditions tested in Fig. 5: PCL2:eGFP alone or normalized by mCherry driven by PCL1, PCL2, PCM1, PCM2, PCH4, or PCH5. Left: eGFP, right: eGFP/mCherry. Each experimental replicate is a unique color. The black line is the CDF for the pooled data of all six experimental replicates. The p-values of one-sample Kolmogorov-Smirnov tests appear to the right of each CDF, colored by experimental replicate. Asterisks indicate p-values: * < 0.05, ** < 0.01, *** < 0.001, and ns = not significant. n=32 per experimental replicate, 6 experimental replicates.

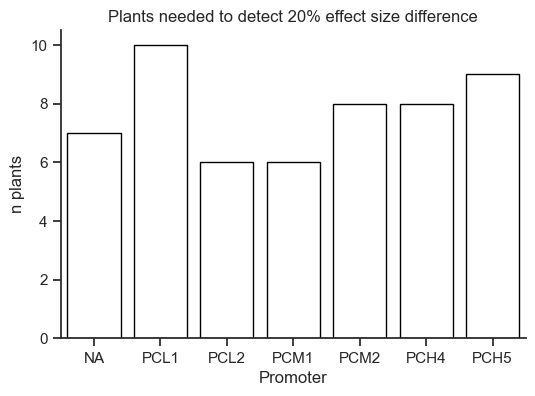

**Supplementary Figure S16.** Number of plants needed to detect a 20% effect size difference given the CVs of the conditions tested in Fig. 5.
